# Topoisomerase II inhibitors induce cGAS-STING dependent inflammation resulting in cytokine induction and immune checkpoint activation

**DOI:** 10.1101/764662

**Authors:** R. D. A Wilkinson, N. McCabe, E. E. Parkes, E.M. Barros, D. I. Johnston, R.M.M Ali, K. Lappin, R.A. Greenberg, D. P. Harkin, S. A. McIntosh, R. D. Kennedy, K. I. Savage

## Abstract

Tumours with genomic instability demonstrate enhanced immunogenicity and potential for response to immune checkpoint blockade (ICB). We previously demonstrated activation of the cGAS-STING pathway following loss of DNA repair, resulting in cytokine induction, lymphocytic infiltration and immune checkpoint activation. Here we explore the role of chemotherapies in inducing this innate immune response, identifying topoisomerase II (topo-II) inhibitors, particularly doxorubicin and epirubicin, as potent inducers of a cGAS-STING dependent interferon response. Mechanistically, topo-II inhibition resulted in significant induction of cytoplasmic DNA and subsequent micronuclei formation, a requirement for efficient cGAS-STING activation and consequent cytokine and immune checkpoint gene induction. Importantly, increased cytokine and immune checkpoint gene expression, as well as increased immune cell infiltration, was also observed in patient derived breast tumour biopsies following topo-II inhibitor-based treatment. Taken together, this study indicates topo-II inhibitors such as doxorubicin, may be best placed to induce immunogenic inflammation, and thereby increase responses to ICB therapies.

**Significance:** This work demonstrates how topo-II inhibitors induce STING-pathway activation, cytokine induction and immune checkpoint protein upregulation in cancer cells and provides a rationale for combining topo-II inhibitors with ICB therapy in early breast cancer.

## Introduction

Defective BRCA/Fanconi anaemia DNA pathway repair machinery has been reported in an estimated 25% of breast cancers [1]. Furthermore, an estimated 60-69% of triple negative breast cancers (tumours lacking oestrogen receptor, progesterone receptor, and HER2 amplification) display a “BRCAness” genomic instability phenotype, with features that mimic BRCA1/2 mutated tumours [2]. Importantly, genomic instability correlates with benefit from ICB, with emerging predictive biomarkers, such as tumour mutational burden, demonstrating promise in the clinic [3].

Response to ICB in genomically unstable tumours has been attributed to accumulation of mutations resulting in neo-antigen production. ICB, such as targeting Programmed cell Death protein 1 (PD-1), then enables an immune response to these neo-antigens through reactivation of tumour-infiltrating lymphocytes. However, we recently reported an additional mechanism for immune activation in double strand break repair (DSBR)-deficient breast cancers, as a result of cytoplasmic DNA originating from defective DNA repair. Damaged cytoplasmic DNA triggers the cGAS-STING pathway, resulting in IRF3 activation and a resultant Type-I interferon response-like transcriptional cascade [4]. Additionally, we have generated a 44-gene expression based assay for identifying activation of this pathway in FFPE tumour samples termed the DNA Damage Immune Response (DDIR) assay, formerly known as the DNA damage response deficiency (DDRD) assay [5, 6]

The majority of breast cancers do not respond to single agent ICB [7]. A potential method of improving response rates may be to enhance the immunogenicity of the tumour. STING agonists, cyclic dinucleotides administered by intra-tumoural injection, are now in clinical trials in combination with ICB. However, as radiotherapy and chemotherapy also result in STING pathway activation [4, 8, 9], the role of these as immune adjuvants is also of interest.

Although the exact mechanisms linking therapeutic DNA damage to cGAS-STING activation remain to be fully elucidated, studies of ionising radiation suggest the formation of micronuclei may be required. Micronuclei are formed when damaged or lagging chromosome fragments are unable to be incorporated into the nucleus during mitosis and are instead independently compartmentalised into cytoplasmic micronuclei [8, 9]. cGAS co-localises with damaged DNA within these micronuclei, which rupture during mitosis, resulting in cGAS-mediated 2’3’ cGAMP production and subsequent STING activation, resulting in IRF3/TBK1 activation and downstream transcriptional induction of Type-I interferon stimulated genes [8, 9].

An extensive number of clinical trials are in progress using combinations of ICB with various chemotherapeutic agents in a number of solid tumours, resulting in mixed outcomes [10]. These studies, however, are largely empirical with the addition of immune checkpoint inhibiting drugs to conventional chemotherapeutic regimens. It is possible, however, that certain chemotherapeutic agents may be more effective than others in the activation of an immune response [11].

To identify chemotherapeutic agents that maximally activate the cGAS-STING pathway, we assessed induction of two key cGAS-STING activated cytokines CXCL10 and CCL5 following treatment with IC_30_ doses of various chemotherapies. Here we show that the topo-II inhibitors doxorubicin and epirubicin are potent activators of the cGAS-STING pathway, which is dependent on micronuclei formation following DNA damage.

## Materials and Methods

### Cell lines, Nocodazole block and generation of chemotherapeutic IC_30_ values

HeLa cells (human cervix adenocarcinoma) (ATCC) were cultured in Dulbecco Modified Eagle Medium containing 10% FCS. MCF10A (non-tumour breast epithelial) were cultured in DMEM/F12 media supplemented with 5% Horse serum, 100 ng/mL Cholera toxin, 10 ng/mL insulin, 20 ng/mL EGF and 0.5 μg/mL hydrocortisone. Cells were maintained in 5 % CO_2_ at 37 °C. All subsequent cells have been derived from original ATCC validated stocks. For G2/M arrest using Nocodazole, cells were incubated in media containing 100 ng/mL Nocodazole (ab120630, Abcam UK) for 24 hours to induce G2/M checkpoint arrest. Following incubation, cells were washed with PBS and fresh media added.

For generation of IC_30_ values, cells were treated with chemotherapeutics for 48 hours before assessing viabliity using Cell Titre Glo reagent (Promega, UK), as per manufacturer’s instructions. Lumiscence was read using a Biosciences BioTek plate reader. Values were analysed using GraphPad Prism (V5.03) and IC_30_ values generated.

### qRT-PCR

RNA was isolated from samples using Ribozol (VWR, USA) according to manufacturer’s instructions. Complementary DNA was generated using First Strand cDNA synthesis kit (Roche, Basel, Switzerland). qRT-PCR evaluation was completed using the following primer sets: CXCL10 forward 5’ GGC CAT CAA GAA TTT ACT GAA AGC A 3’ and reverse 5’ TCT GTG TGG TCC ATC CTT GGA A 3’, CCL5 forward 5’ TGC CCA CAT CAA GGA GTA TTT 3’ and reverse 5’ CTT TCG GGT GAC AAA GAC G 3’ and PUM1 forward 5’ CCA GAA AGC TCT TGA GTT TAT TCC 3’ and reverse 5’ CAT CTA GTT CCC GAA CCA TCT C 3’. Differences in expression were detected using a Roche LightCycler^®^480.

### Western blotting

Western blotting was carried out as previously described [4] using the following antibodies/dilutions; anti-γH2AX (1/2000; JBW301-MerckMillipore), anti-β-actin (1/10000; A2228-Sigma), anti-STING (1/1000, 13647-Cell Signalling), anti-cGAS (1/1000; 15102-Cell Signalling), anti-vinculin (1/1000; 14-9777-82-Thermo), anti-mouse-HRP (1/10000; 7076-Cell signalling), anti-rabbit-HRP (1/10000; 7074-Cell Signalling).

### Immunofluorescence

Immunofluorescent staining was carried out as previously described [4] using the following antibodies/dulitions; anti-dsDNA (1/50; Santa Cruz Biotechnology), anti-γH2AX (1/2000; JBW301-MerckMillipore), anti-Lamin B1 (1/1000; ab16048-Abcam), anti-mouse-IgG AlexaFlour-488 (1/1000; ab150113-Abcam), anti-rabbit-IgG AlexaFlour-594 (1/1000; A-11072-ThemoFischer), anti-mouse-IgG AlexaFlour-594 (1:1000; ReadyProbes). Nuclear DNA was stained using Hoechst-33342 (5 μg/mL-SIGMA).

### siRNA reverse transfection

Cells were reverse transfected with siRNA (10 nM) using Lipofectamine RNAiMAX (Life Technologies, UK) using the following siRNAs; siSTING_#1: 5’-CAGCGGCUGUAUAUUCUCCUCCC-3’, siSTING_#2: 5’-GGUCAUAUUACAUCGGAUAUU-3’, sicGAS_#1: 5’-AGAGAAAUGUUGCAGGAAAUU-3’, sicGAS_#2: 5’-CAGCUUCUAAGAUGCUGUCAAAG-3’. Cells were collected for analysis 72-hours post-transfection.

### Patient samples

Pre-and on-treatment neoadjuvant chemotherapy breast tumour samples were collected by 14G ultrasound guided needle core biopsy (Ethical approval obtained from the Office of the Research Ethics Committee Northern Ireland, reference 13/NI/0107).

## Results

### Inhibition of topoisomerase I and II strongly induces CXCL10/CCL5 expression

In order to identify drugs that could activate the cGAS-STING pathway, we selected a variety of compounds representative of classes of chemotherapy used in the clinic. These included the intrastrand/interstrand crosslinking agents cisplatin and mitomycin C, the anti-mitotic agents paclitaxel and vinorelbine, the topo-I inhibitor irinotecan, the topo-II inhibitors doxorubicin and etoposide and the anti-metabolite 5-fluorouracil. In order to allow the evaluation of immune activation at equivalent cytotoxic concentrations, 48 hour IC_30_ values were identified in HeLa cells **(Table S1).** CXCL10 and CCL5 expression, two STING-dependent cytokines associated with lymphocyte recruitment to the tumour microenvironment, was then assessed by qRT-PCR. With the exception of the anti-mitotics, we observed a significant increase in CXCL10 and CCL5 expression with all compounds versus the vehicle treated control, with the greatest increases observed for irinotecan (11.6- and 32.8-fold), doxorubicin (74.2- and 52.2-fold) and etoposide (42.7- and 27.8-fold) **(Figure 1A)**. Although all agents, excepting the anti-mitotic agents, induced DNA damage **(Figure S1A)**, activation of CXCL10 and CCL5 did not appear to correlate with levels of DNA damage as assessed by H2AX phosphorylation (γH2AX) **(Figure S1B).**

**Figure 1.**
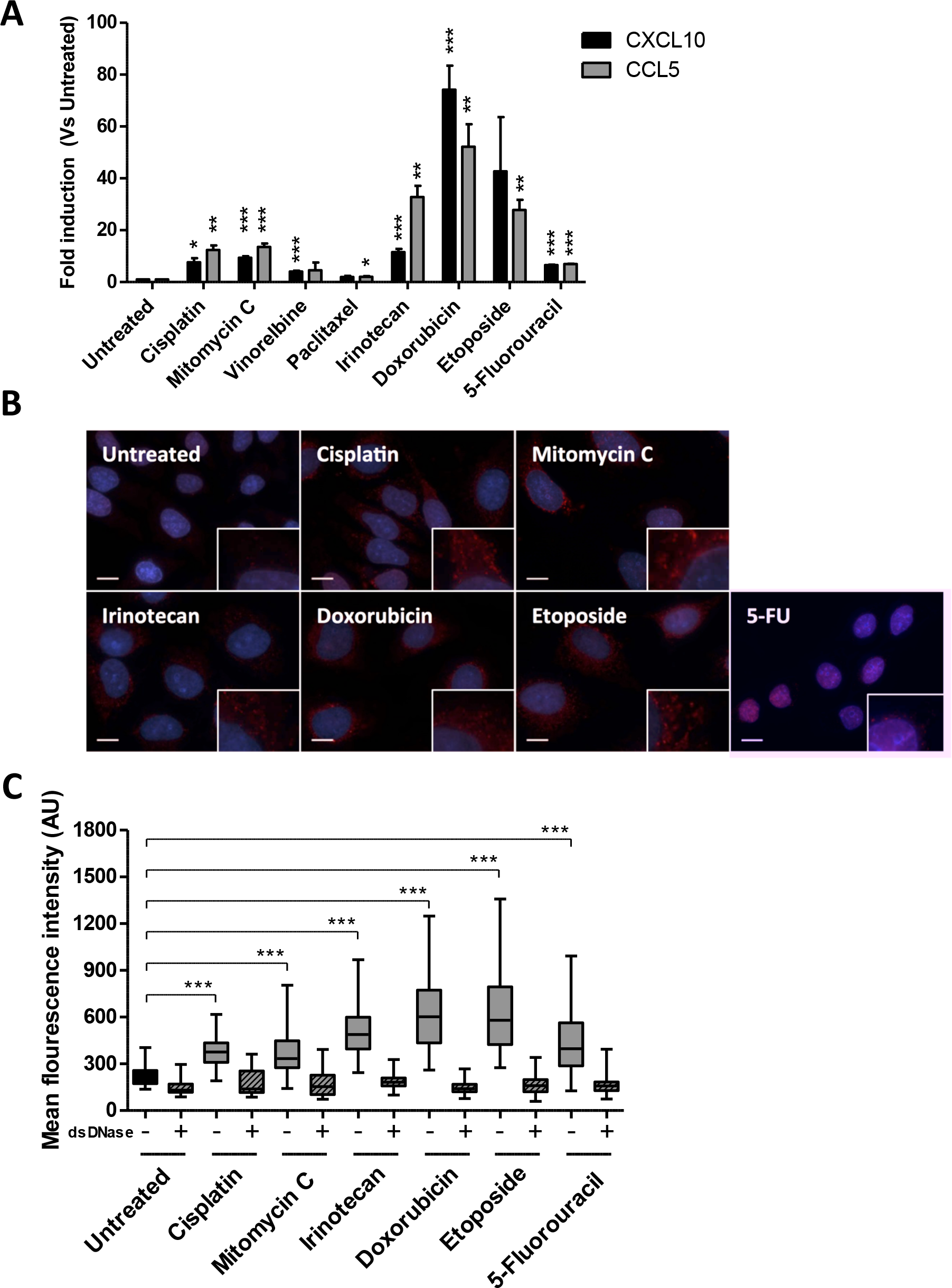
Inhibition of topoisomerase I and II causes the greatest increase in CXCL10/CCL5 expression in HeLa cells. A) Treatment with 48 hour IC_30_ values of chemotherapy significantly increased CXCL10 and CCL5 expression in HeLa cells. B) Treatment with 48 hour IC_30_ values of DNA damaging chemotherapeutics causes an increased translocation of dsDNA into the cytosol in HeLa cells. C) Quantification of cytosolic dsDNA shown in B. dsDNAse treatment was used as a control to confirm quantified fluorescence signal was dsDNA (***=p≤0.001).

cGAS is a cytoplasmic protein and is known to be activated by cytosolic double stranded DNA (dsDNA), after which it catalyses the production of 2,3-cGAMP, which in turn activates STING [12]. We therefore assessed cytosolic dsDNA following chemotherapy treatment using immunofluorescent staining with a dsDNA specific antibody. Treatment with cisplatin, mitomycin C, irinotecan, doxorubicin, etoposide and 5-FU, all resulted in cytosolic dsDNA **(Figure 1B).** Quantification using a previously described image compartmentalisation and quantification algorithm [13], confirmed that cytosolic dsDNA significantly increased following treatment with cisplatin, mitomycin C, irinotecan, doxorubicin, etoposide and 5-FU, in comparison to vehicle treated control cells **(Figure 1C)**.

### Topoisomerase mediated cytokine induction is cGAS-STING-dependent

We next assessed the dependency of CXLC10 and CCL5 expression on the cGAS-STING pathway. Depletion of either cGAS or STING (using 2 independent siRNAs) 48-hours prior to treatment with cisplatin, mitomycin C, irinotecan, doxorubicin or etoposide led to a significant attenuation in CXCL10 and CCL5 induction **(Figure 2A-B & S2A-B)**. As doxorubicin treatment consistently resulted in the greatest increase in CXCL10 and CCL5 expression, we focused our subsequent analysis on this compound. Similarly, doxorubicin treatment of isogenic CRISPR knockout MCF10A cells (normal-like breast cells) harbouring loss of cGAS (crcGAS) or STING (crSTING) resulted in a 12.5- and 10.5-fold increase in CXCL10 and CCL5 expression in control cells, with significant attenuation of CXCL10 and CCL5 induction in cGAS and STING null cells **(Figure 2C & S2C-D)**. Importantly, induction of cytosolic dsDNA was independent of cGAS and STING **(Figure S2E).** Together these data indicate that the cytosolic DNA, as a result of DNA damage, is upstream of cGAS-STING and that cytokine expression is dependent on cGAS and STING.

**Figure 2.**
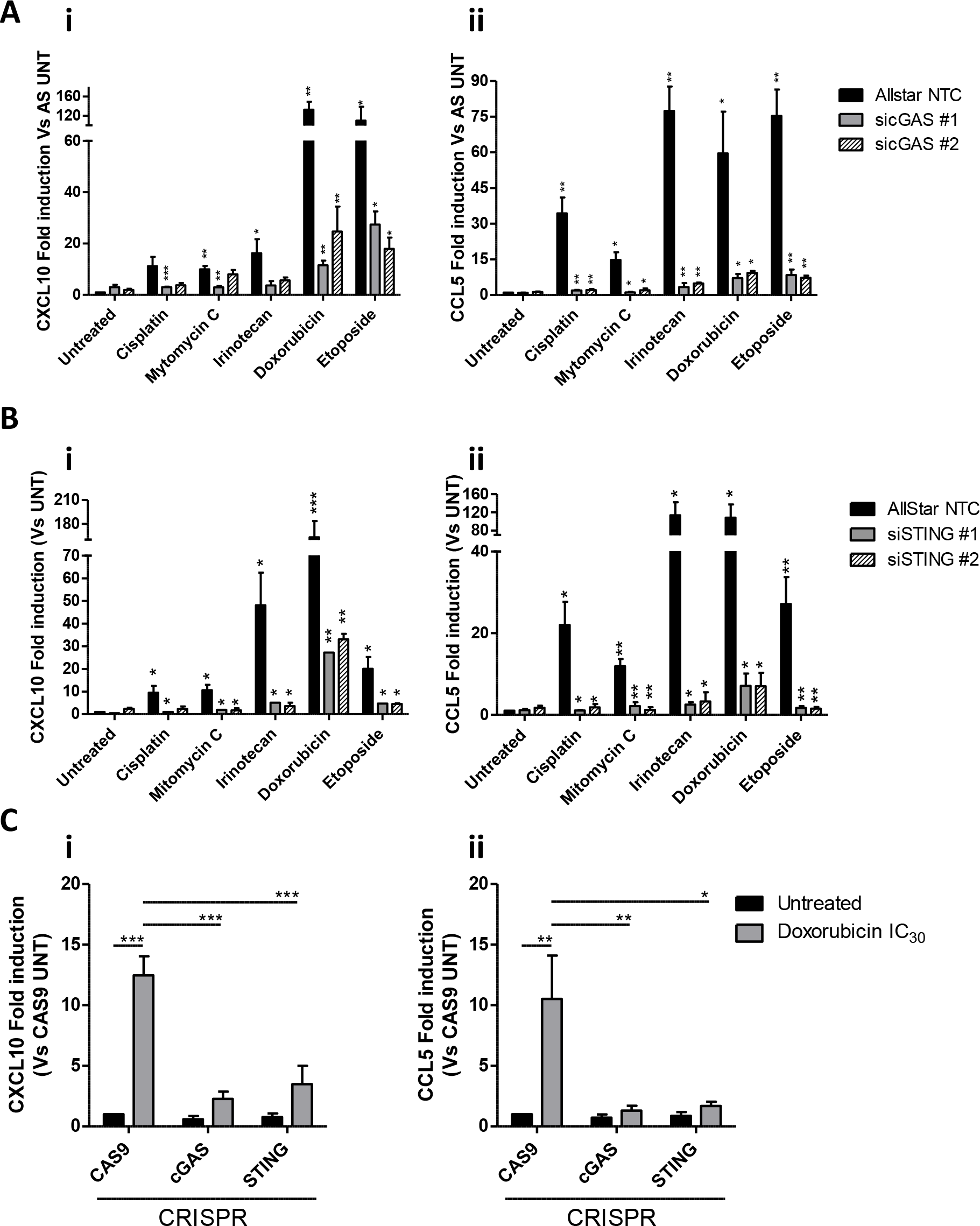
Chemotherapeutic treatment stimulates CXCL10 and CCL5 in a cGAS and STING dependent manner. qRT-PCR analysis of CXCL10 and CCL5 in control and (A) cGAS and (B) STING depleted HeLA cells, following 48-hour treatment with indicated drugs C) qRT-PCR analysis of CXCL10 and CCL5 expression in MCF10A cells with CRISPR knock out of cGAS or STING (control cells express Cas9 only, with no gene targeting guide RNA), following 48-hour treatment with IC_30_ doxorubicin. All data represents mean of 3 independent experiments +/− SEM (*=p≤0.05, **=p≤0.01, ***=p≤0.001).

### Treatment with doxorubicin activates CXCL10/CCL5 expression in a micronuclei dependent manner

Previous reports have indicated that cGAS concentrates within radiation-induced micronuclei, which contain broken DNA fragments formed due to unrepaired DSBs that persist through mitosis. These structures are enveloped by a micronuclear membrane, which ruptures during mitotic progression releasing DNA and activated cGAS into the cytoplasm, resulting in potent STING activation and a subsequent Type I interferon response [8, 9]. We therefore asked if this mechanism also underpinned the activation of cGAS-STING observed in response to doxorubicin. First we evaluated the expression of CXCL10 and CCL5 over a 48-hour time-course in HeLa cells treated with an IC_30_ dose of doxorubicin. We observed small increases in CXCL10 and CCL5 expression at early time points following doxorubicin treatment (5-12 fold) up to 24 hours post-treatment, with marked increase in CXCL10 (50.1-fold increase) and CCL5 (53.3-fold increase) expression 48-hours post-treatment **(Figure 3A).** Using immunofluorescent imaging we also assessed DNA damage and micronuclei formation using antibodies targeting γH2AX and Lamin B1 **(Figure 3B).** We observed a peak in DNA damage and micronuclei positive cells 24-hours after doxorubicin treatment **(Figure 3C-D).** Together, these data indicate that doxorubicin induces micronuclei that are temporally associated with cytokine induction. To further explore this, we blocked micronuclei formation in doxorubicin treated cells by co-treating with Nocodazole, a microtubule depolymerising agent that results in early mitotic arrest, and assessed CXCL10 and CCL5 induction **(Figure S3A-B).** This resulted in complete abrogation of doxorubicin induced CXCL10 and CCL5 induction **(Figure 3E & S3C).** Importantly, doxorubicin induced DNA damage was unaffected by nocodazole treatment, despite a complete loss of micronuclei induction in these cells **(Figure 3F-G & S3D-E).** Taken together these data suggest that S-phase DNA damage results in micronuclei formation during mitosis that release cytosolic DNA and induce cGAS-STING dependent cytokine expression. Importantly, this data also suggests that anti-mitotic agents, e.g. taxanes, when used at doses that induce mitotic arrest, may block cGAS-STING activation through inhibition of mitotic progression and subsequent micronuclei formation and/or rupture. To test this, we also assessed the effect of combined paclitaxel and doxorubicin treatment on micronuclei formation and CXCL10/CCL5 induction. Indeed, the doxorubicin induced upregulation of CXCL10 and CCL5 was significantly abrogated by co-treatment with IC_30_ doses of paclitaxel **(Figure 3H-I & S3D-E)**. Importantly, the dose of paclitaxel used in these experiments did not affect doxorubicin induced DNA damage but did result in mitotic arrest and blocked doxorubicin induced micronuclei formation **(Figure S3F-H).**

**Figure 3.**
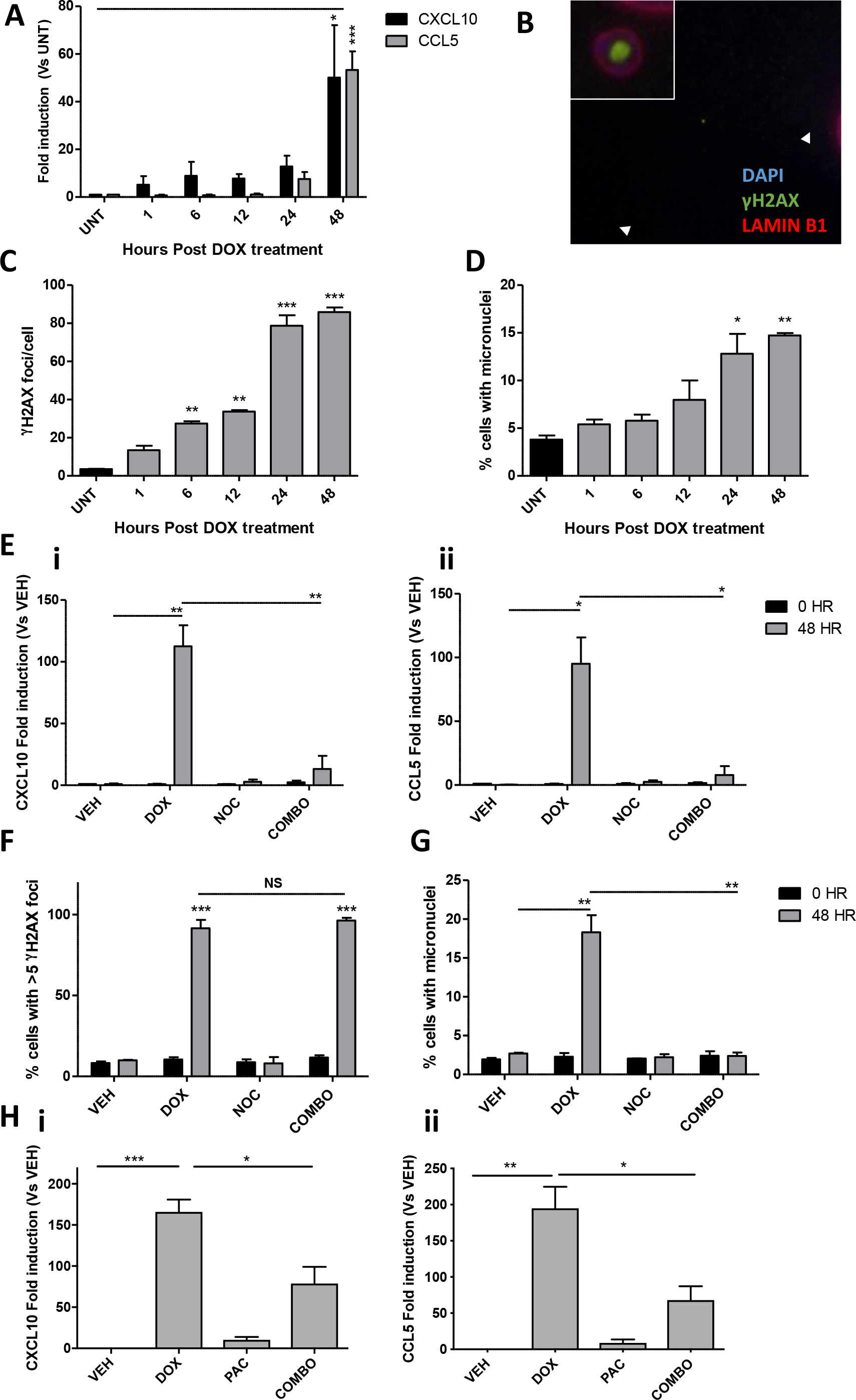
Doxorubicin induced CXCL10/CCL5 expression is micronuclei dependent. A) Time course of CXCL10/CCL5 expression in HeLa cells following treatment with IC_30_ doxorubicin. B) Representative image of immunofluorescent staining of γH2AX and LAMINB1 following 48 hours doxorubicin treatment. C) Quantification of γH2AX foci in HeLa cells over a 48 hour time course treatment as shown in B. D) Percentage of cells with micronuclei evaluated at the indicated time points following treatment. E) qRT-PCR mediated quantification of CXCL10 (i) and CCL5 (ii) in HeLa cells treated with vehicle, doxorubicin (Dox), Nocodazole (Noc), or combined Dox and Noc (Combo) for the indicated timepoints. F) Confirmation of doxorubicin induced DNA damage in HeLa cells treated as in E. G) Percentage of cells shown in E with micronuclei evaluated at the indicated timepoints. H) qRT-PCR mediated quantification of CXCL10 (i) and CCL5 (ii) in HeLa cells treated with vehicle, doxorubicin (Dox), Paclitaxel (Pac), or combined Dox and Pac (Combo). All quantitative data represents mean of 3 independent experiments +/− SEM (*=p≤0.05, **=p≤0.01, ***=p≤0.001).

### Topo-II inhibition induces cytokine and immune checkpoint expression in cell line models and breast tumours

Both doxorubicin and epirubicin are used in the neoadjuvant and adjuvant treatment of early breast cancer. Therefore, we confirmed if epirubicin had a similar effect to doxorubicin *in vitro* in terms of induction of cytokine expression. CXCL10 was induced between 120- and 88-fold and CCL5 induced between 118- and 110-fold following chemotherapy **(Figure 4A).** Epirubicin also induced similar levels of DNA damage, cytosolic DNA and micronucleated cells **(Figure S4A-C)**. We and others have previously reported that activation of the cGAS-STING pathway not only drives cytokine induction and tumour lymphocytic infiltration, but also leads to transcriptional upregulation of immune checkpoint genes including PD-L1, PD-L2, GAL9 and HVEM [4]. Therefore, we investigated the effect of doxorubicin and epirubicin treatment on the expression of these well established immune checkpoint genes. Indeed, expression of all of these genes was significantly upregulated following treatment with either doxorubicin or epirubicin **(Figure 4B & S4D-E).** We also carried out RNA-Seq analysis and examined DDIR signature scores in HeLa cells following treatment with doxorubicin or epirubicin. This showed a significant increase in DDIR scores, as well as induction of CXCL10 and CCL5, following both doxorubicin and epirubicin treatment **(Figure 4C)**. Examining this data further revealed induction of a panel of classical Type I interferon response genes, as well as immune checkpoint genes, following treatment with either doxorubicin or epirubicin **(Figure 4C)**. This suggests that topo-II inhibition leads to activation of the cGAS-STING pathway resulting in increased immune checkpoint gene expression in addition to cytokine expression.

**Figure 4.**
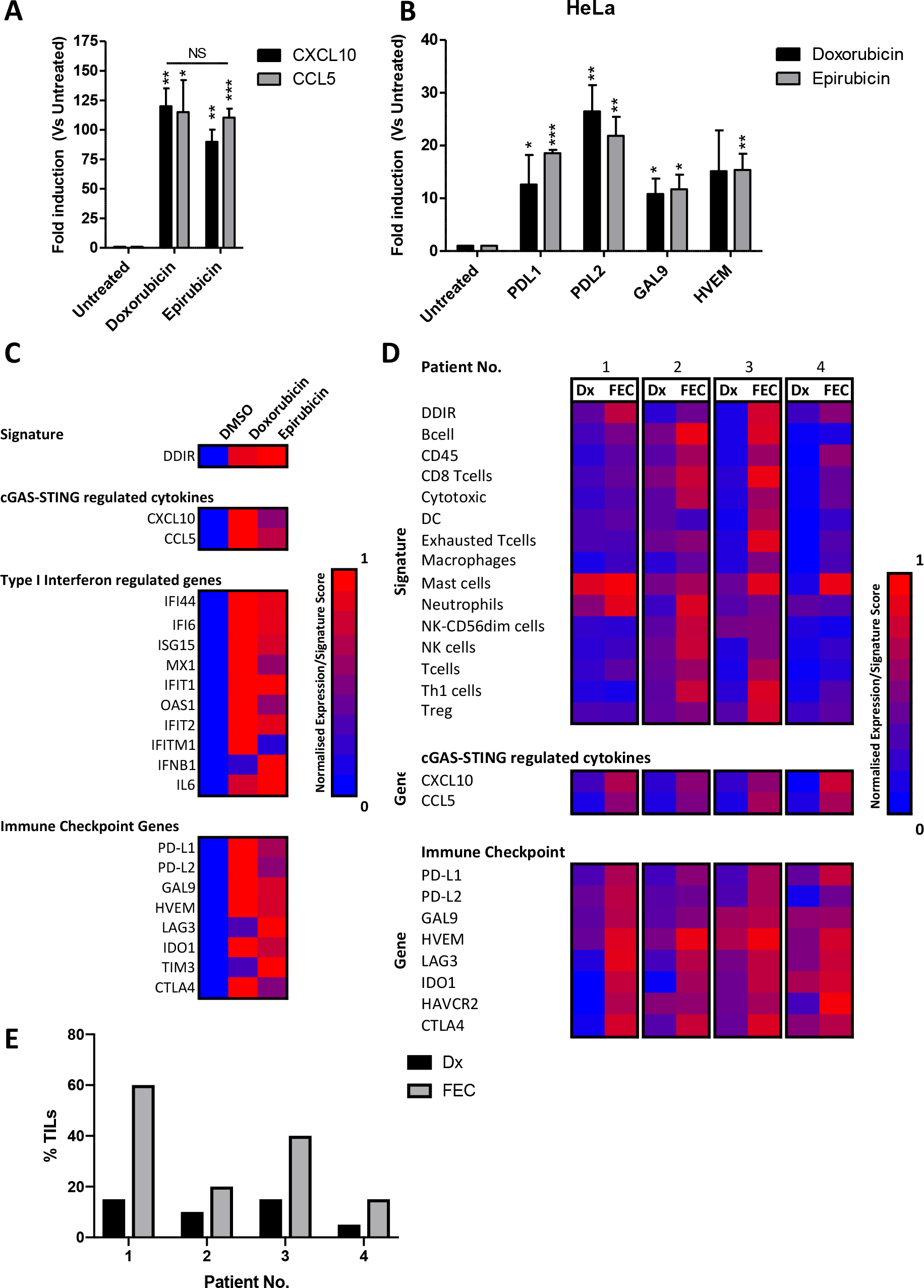
Anthracycline treatment increases CXCL10 and CCL5 expression using patient samples. qRT-PCR analysis of A) CXCL10 and CCL5 and B) Immune checkpoint gene expression following 48 hours treatment with IC_30_ doxorubicin or epirubicin in HeLa cells. All data represents mean of 3 independent experiments +/− SEM (*=p≤0.05, **=p≤0.01, ***=p≤0.001). C) RNA-Seq generated DDIR signature and individual gene expression values from vehicle treated (DMSO) HeLa cells and following treatment IC_30_ doses of doxorubicin or epirubicin for 48hrs. D) RNA-Seq generated gene signature scores (top Panel) and individual gene expression values from pre-treatment diagnostic (Dx) and post 3-cycles of FEC treatment (FEC) breast tumour core biopsies from 4 individual patients (labelled 1-4). E) % TILs quantified H+E stained sections from the same core biopsies utilised for RNA-Seq analysis shown in D.

We hypothesised that *in vivo*, this would lead to tumour lymphocytic infiltration, but limited immune mediated tumour killing. To test this, we performed RNA-seq analysis of breast tumour core biopsies from 4 early breast cancer patients at diagnosis and after 3-cycles of neoadjuvant FEC (5-fluorouracil, epirubicin, cyclophosphamide) chemotherapy. These patients were chosen as they did not respond to neoadjuvant therapy (assessed via Residual Cancer Burden scoring following tumour resection), suggesting that none of these tumours had an intrinsic DNA repair defect and were immunogenically “cold” at diagnosis [14]. Consistent with this, all four patients were DDIR negative at diagnosis **(Figure 4D).** In keeping with our pre-clinical data, we observed an increase in expression in CXCL10 and CCL5 following 3-cycles of FEC chemotherapy **(Figure 4D).** Additionally, all 4 tumours converted from DDIR negative at diagnosis, to DDIR positive following treatment **(Figure 4D).** Moreover, expression of many well characterised immune checkpoint genes, including the clinically targeted PD-L1 and CTLA4 genes, increased in all patients’ tumours following FEC therapy **(Figure 4D).** We next studied the presence of tumour infiltrating lymphocytes (TILs) following doxorubicin treatment using H+E staining of tumour sections. As expected, all four demonstrated increased TILs post-chemotherapy **(Figure 4E).**Well established gene expression signatures were used to characterise the TILs further [15]. Consistent with the histology, signature scores for the majority of immune cells, particularly T-cells and cytotoxic T-cells, increased in these tumours following FEC treatment **(Figure 4D)**. Importantly, we observed increased signature scores for exhausted T-cells following chemotherapy treatment, in keeping with the expression of immune checkpoint genes in these tumours **(Figure 4D).**

## Discussion

Numerous studies have now shown that DNA repair pathway deficiencies can stimulate inflammatory cascades, and examples include loss of core DNA repair proteins including NSB1, CHK2, BRCA1, FANCD2, ATM, MRE11, [16] [4], [16, 17], [18]. Furthermore, inhibition of specific DNA repair proteins (such as PARP1) has also been linked with immune activation [19]. In keeping with this, a number of groups have demonstrated stimulation of type 1 interferons and cytokines following treatment with DNA damaging agents or ionising radiation in both the *in vitro* and *in vivo* settings [4, 8, 9, 20]. Similarly, a recent study screening a panel of FDA-approved drugs for agents that induce immunogenic cell death (ICD), identified the topo-II inhibitor teniposide as a potent ICD inducer [21]. ICD is a mode of cell death reported to enhance release of tumour specific antigens from dying cells and thereby boost immune cell recruitment and T-cell mediated cancer cell death. This study also went on to show that *in vitro*, teniposide activates the cGAS/STING pathway leading to a Type-I interferon response [21]. However, teniposide is a poorly tolerated drug, rarely used in the clinic due to toxicity. In contrast, to the best of our knowledge, this is the first time several commonly clinically used chemotherapeutic agents have been compared, in terms of cGAS-STING activation.

In this study, we identified topo-II inhibitors/anthracyclines as potent activators of the cGAS-STING-dependent cytokine response. We found that topo-II inhibitors damage DNA, resulting in the formation of micronuclei and release of dsDNA into the cytosol. This drives expression of inflammatory cytokines CXCL10 and CCL5 in a cGAS and STING dependent manner.

Assessing activation of these cytokines over time revealed a modest induction of both CXCL10 and CCL5 at early time points following doxorubicin treatment. However, we observed a dramatic increase in their induction 48 hours following treatment, which was abrogated by inhibition of mitosis using nocodazole. Micronuclei are formed during mitosis, when lagging dsDNA fragments, produced by DNA damage and/or inefficient repair, are packaged within a distinct nuclear membrane. Previous studies have demonstrated that rupture of ionising radiation induced micronuclei leads to release of dsDNA and activated cGAS, with subsequent potent activation of STING, and a downstream type I interferon response [8, 9]. Our data suggest the same mechanism for cGAS-STING activation following topo-II inhibition.

Given that the cGAS-STING driven immune response has been shown to lead to upregulated expression of immune checkpoint genes, we hypothesised that in patients, treatment with topo-II inhibitors would lead to cGAS-STING dependent transcriptional activation of cytokines and immune checkpoint genes. This promotes immune cell recruitment and tumour infiltration, but prevents immune mediated tumour cell killing via immune checkpoint activation. Indeed, analysis of breast tumour biopsy samples collected from patients pre- and post- FEC treatment confirmed these *in vitro* findings, demonstrating increased expression of these key cGAS-STING driven cytokines, and immune checkpoint genes. Additionally, FEC treatment led to increased TILs within all biopsy samples and, using well-validated signatures of different immune cell subtypes, we found that FEC treatment lead to increased scores for many immune cell types, particularly CD8-positive/cytotoxic T-cells. In keeping with our preclinical data, we also observed upregulation of key immune checkpoint genes, including PD-L1, in all breast tumours following FEC treatment.

Although trials have been conducted using single agent immune checkpoint inhibitors in breast cancer, recent efforts have focused on identifying effective combination therapies. These have included empirically combining well-characterised and commonly used chemotherapeutic classes (such as anti-metabolites, anti-microtubule and crosslinking agents) with novel ICB treatments to improve outcome [10]. Our data suggests there may be a rationale for combining topo-II inhibitors with ICB agents. Importantly, the recently reported TONIC trial aimed to utilise different modes of DNA damage to enhance immune responses to the anti-PD1 agent nivolumab in metastatic TNBC. In this study, patients received no induction therapy, or induction with hypofractionated radiotherapy, cyclophosphamide, cisplatin or doxorubicin, followed by 3 cycles of nivolumab. The highest response rate (35%) was seen in patients receiving induction with doxorubicin [20]. Furthermore, patients pre-treated with doxorubicin showed increased tumoural expression of immune checkpoint genes PD1 and PDL1 [20]. Crucially, scheduling of these treatment regimens is of paramount importance. Patients in the TONIC trial were given two, low dose (15mg) treatments of doxorubicin, followed by nivolumab treatment. However, it is not clear whether this dose was optimal for immune priming, and/or whether maintenance of doxorubicin during nivolumab therapy is needed in order to maintain immune activation and thereby enhance response to ICB therapy. The Gepar Nuevo trial has also investigated the potential of immunotherapy combinations in the neoadjuvant setting in early breast cancer. Patients with TNBC were treated with either durvalumab (anti-PD-L1) or placebo, combined with 4 cycles of nab-paclitaxel followed by epirubicin/cyclophosphamide. A higher pathological complete response rate was seen in ICB combination patients, compared with chemotherapy alone (53.4% versus 44.2%) [22]. Despite this trial not having a pre-ICB immune induction arm, it raises the question whether induction with anthracycline chemotherapy before subsequent combination, and/or maintenance during treatment, will be the most effective therapeutic approach.

Finally, another empirical ICB combination trial, IMPASSION-130, showed improved progression-free survival following nab-paclitaxel in combination with atezoluzimab, compared with nab-paclitaxel alone, in metastatic triple negative breast cancer [23]. This led to the Federal Drug Administration’s approval of atezoluzimab (targeting PD-L1) in combination with nab-paclitaxel in the treatment of PD-L1 positive advanced triple negative breast cancer. However, our pre-clinical data presented above suggests that taxanes may in fact supress cGAS-STING activation, via blocking formation of cGAS-activating micronuclei. This suggests that taxanes may not be the ideal class of drugs for immune priming in combination with ICB agents. Intriguingly, although IMPASSION-130 evaluated nab-paclitaxel in combination with atezoluzimab, the investigators did not assess the efficacy of atezoluzimab alone, which may have demonstrated equivalent or better efficacy than the treatment combination.

In conclusion, our data shows that commonly-used topo-II inhibitors potently activate the cGAS-STING innate immune pathway. We have demonstrated that the underlying mechanism of cGAS-STING activation is through the induction of cytoplasmic dsDNA and micronuclei formation. Thus, these agents may have a clinical role in converting immune “cold” tumours to “hot” tumours and may suggest a biological rationale for logical combinations with immune checkpoint therapies. This is supported by data from the TONIC trial; nevertheless, further clinical studies to optimise dose and scheduling of combination therapies are required. Additionally, the use of biomarkers, such as DDIR, to identify immunogenically cold tumours need to be clinically validated and utilised to identify patients whom will benefit from these combinations.

