## Supplementary Table & Figures for "Topoisomerase II inhibitors induce cGAS-STING dependent inflammation resulting in cytokine induction and immune checkpoint activation"

### Supplementary Data

| | HeLa | | Mechanism of action |
| --- | --- | --- | --- |
| **Units in µM** | $IC_{30}$ | $IC_{50}$ | |
| Cisplatin | 12.75 | 15.79 | Crosslinking agent |
| Mitomycin C | 3.89 | 5.55 | Interstrand crosslinking agent |
| Paclitaxel | 0.0029 | 0.004 | Targets mitotic spindle assembly, blocking mitosis, activating cell checkpoint signalling |
| Vinorelbine | 0.0104 | 0.0137 | Targets mitotic spindle assembly, blocking mitosis, activating cell checkpoint signalling |
| Irinotecan | 45.28 | 97.29 | Metabolised into SN-38, an inhibitor of topoisomerase I |
| Doxorubicin | 0.3 | 0.8757 | Doxorubicin interacts with DNA by intercalation, disrupting topoisomerase II activity |
| Etoposide | 10.58 | 19.82 | Forms complex between DNA and Topoisomerase II enzyme, and prevents ligation of DNA strands |
| 5-Fluorouracil | 1.11 | 1.98 | Antimetabolite, disrupts action of thymidylate synthase stopping production of new DNA |

**Supplementary Table 1:** Mechanism of action and $IC_{30}$ and $IC_{50}$ values of the indicated

A

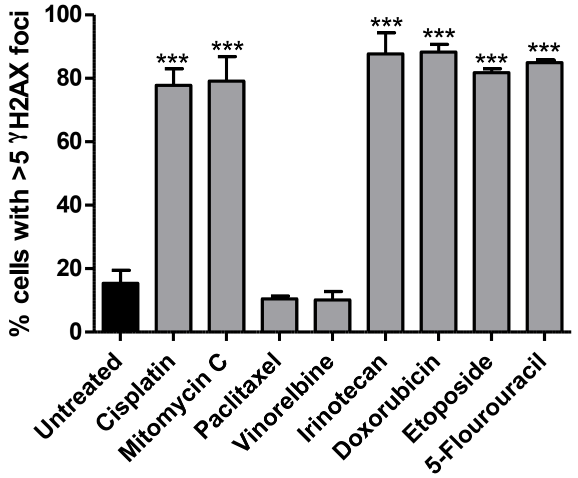


B

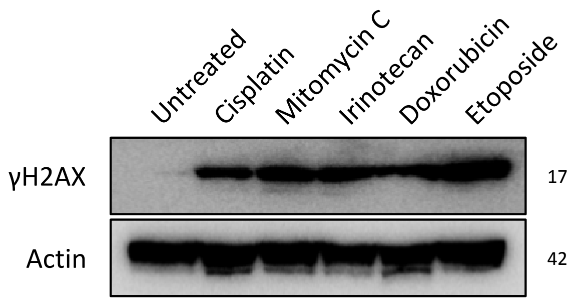


**Supplementary Figure 1.** A) Quantification of γH2AX foci positive (>5 foci) cells following treatment of HeLa cells with 48-Hour IC30 values of the indicated chemotherapeutics B) Representative Western blot analysis of γH2AX levels in HeLa cells with 48-Hour IC30 values of the indicated chemotherapeutics.

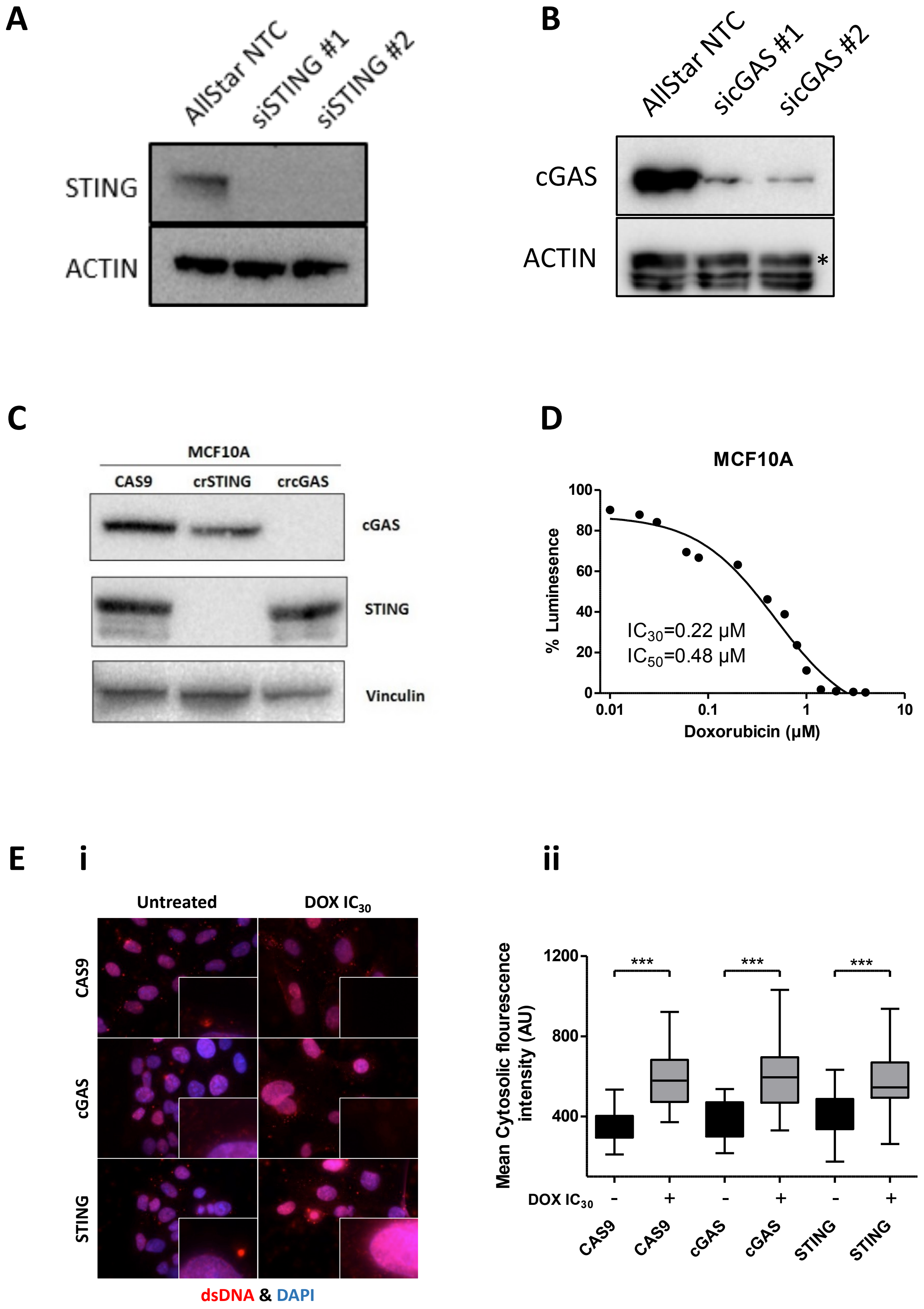


**Supplementary Figure 2.** A-B) Representative Western blots demonstrating typical depletion of STING (A) and cGAS (B) with the indicated siRNAs in HeLa cells. C)

Representative Western blots demonstrating depletion of cGAS and STING in CRISPR edited MCF10A cells. Control cells express Cas9 but no gene targeting guide RNA. D) 48 hour dose response of MCF10A cells with doxorubicin. Data represents mean response of three independent experiments. E) 9i) Representative images of immunofluorescent dsDNA staining in MCF10 cells, including cGAS and STING CRISPR knockout lines. (ii) Quantification of cytoplasmic dsDNA from three independent experiments represented in (Ei). Data represents mean cytoplasmic dsDNA fluorescence intensity from three independent experiments (***=$p \leq 0.001$).

### Supplementary Figure 3

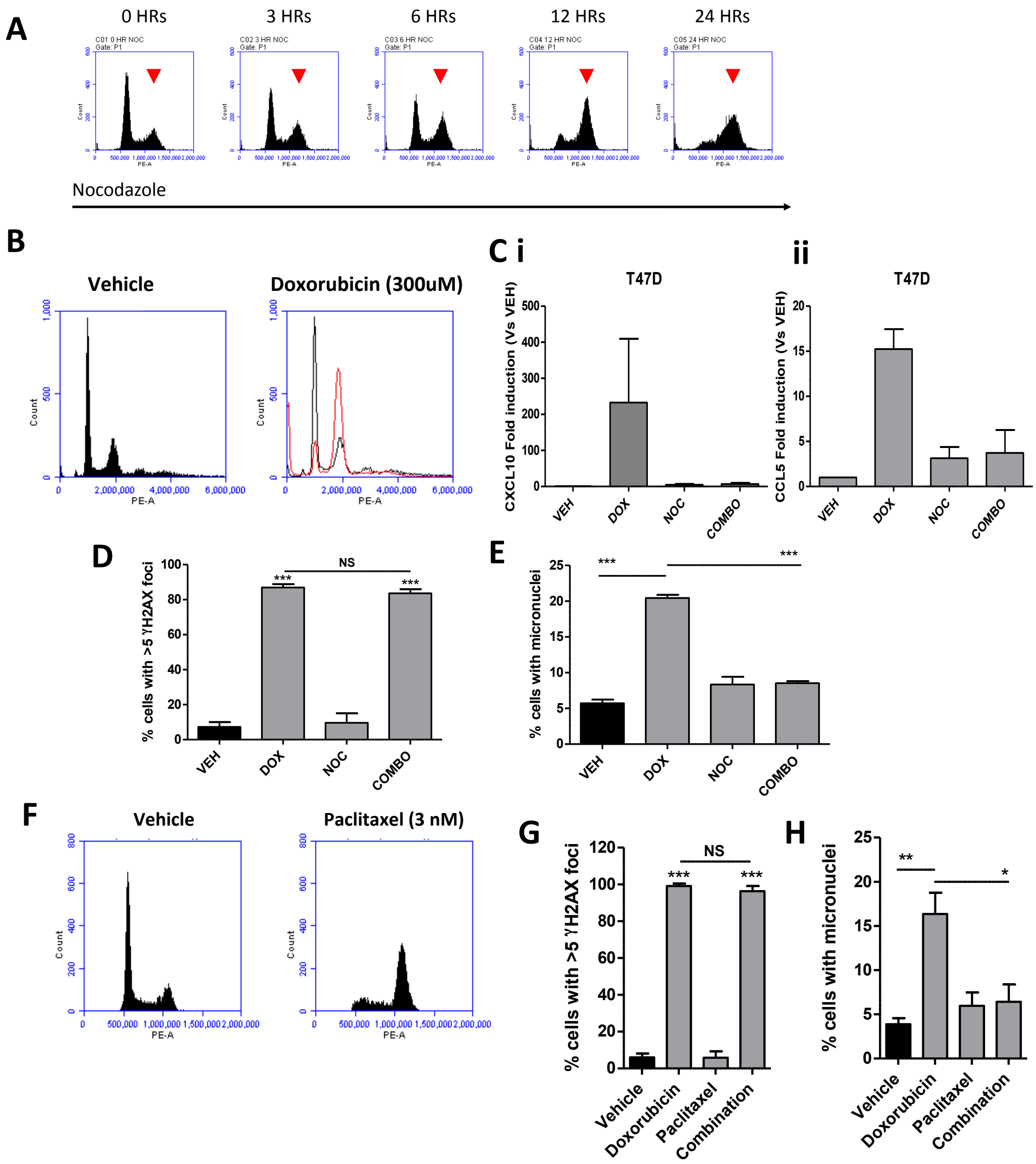


**Supplementary figure 3.** A) Representative FACs profiles of PI stained HeLa cells following Nocodazole treatment for the indicated time-points, demonstrating efficient G2/M block after which doxorubicin was added to cells for 48 hours to generate data shown in figure 3  B) Representative FACs profiles of PI stained T47D cells following Nocodazole treatment for 24 hours, after which doxorubicin was added to generate data below, demonstrating efficient G2/M block. C) qRT-PCR mediated quantification of CXCL10 (i) and CCL5 (ii) in T47D cells treated with vehicle, doxorubicin (Dox), nocodazole (Noc), or combined dox and noc (Combo) for 48hrs. D) Confirmation of doxorubicin induced DNA damage in T47D cells treated as in C. E) Percentage of cells shown in C with micronuclei evaluated at 48 hrs timepoint. F) Representative FACs profiles of PI stained HeLa cells following Paclitaxel treatment, demonstrating efficient G2/M block after which doxorubicin was added to cells for 48 hours to generate data shown in figure 3. G) Confirmation of paclitaxel induced DNA damage in HeLa cells treated as in Figure 3H. H) Percentage of cells shown in Figure 3H with micronuclei evaluated at 48 hrs following Doxorubicin treatment. All quantitative data represents mean of 3 independent experiments +/- SEM (*=$p \leq 0.05$, **=$p \leq 0.01$, ***=$p \leq 0.001$).

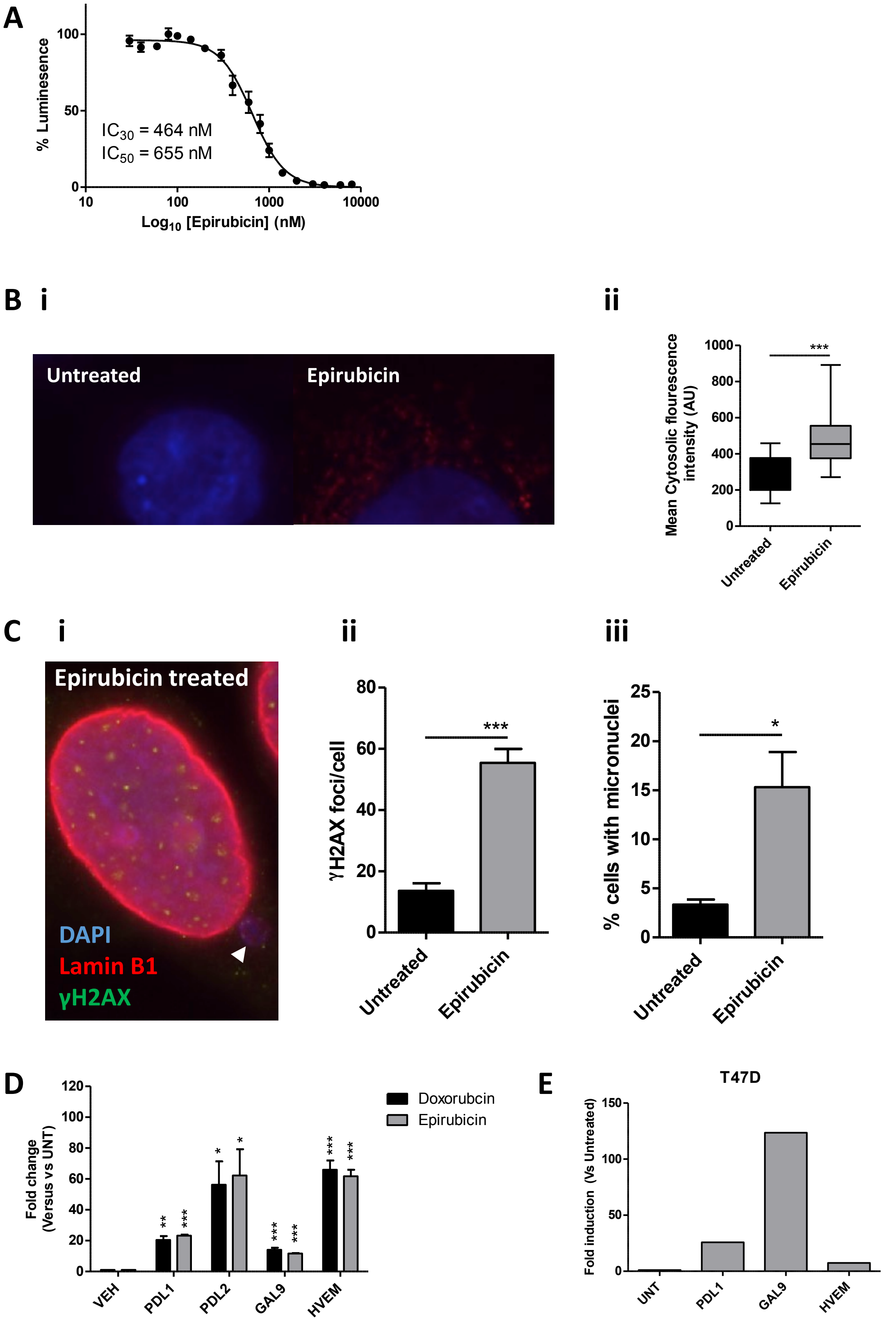


**Supplementary Figure 4.** A) 48 hour dose response in HeLa cells with epirubicin. Data represents mean response of three independent experiments. B) (i).

Representative image of dsDNA staining in HeLa cells following 48-hr treatment with $IC_{30}$ epirubicin. (ii). Quantification of cytoplasmic dsDNA from three independent experiments represented i. C) i. Representative image of γH2AX and Lamin B1 staining in epirubicin treated HeLa cell displaying a micronucleus. ii-iii. Quantification of γH2AX and micronuclei positive cells following 48-hrs treatment with $IC_{30}$ epirubicin. D) Immune checkpoint gene expression following 48 hours treatment with $IC_{30}$ doxorubicin or epirubicin in MCF10A cells E) Immune checkpoint gene expression following 48 hours treatment with $IC_{30}$ epirubicin in T47D cells. PDL2 is not expressed in T47D cells and is therefore not shown. Data represents mean of 3 independent experiments +/- SEM (*=$p \leq 0.05$, **=$p \leq 0.01$, ***=$p \leq 0.001$).
